# An automated, machine learning-based detection algorithm for spike-wave discharges (SWDs) in a mouse model of absence epilepsy

**DOI:** 10.1101/309146

**Authors:** Jesse A. Pfammatter, Rama K. Maganti, Mathew V. Jones

**Author notes:** **Corresponding Author:** Jesse Pfammatter, 5505 WIMR II, 1111 Highland Ave., Madison WI, 53705.

## Abstract

**Objective:** Manual detection of spike-wave discharges (SWDs) from EEG records is time intensive, costly, and subject to inconsistencies/biases. Additionally, manual scoring often omits information on SWD confidence/intensity which may be important for the investigation of mechanistic-based research questions. Our objective is to develop an automated method for the detection of SWDs in a mouse model of absence epilepsy that is focused on the characteristics of human scoring of pre-selected events to establish a confidence-based, continuous-valued scoring.

**Methods:** We develop a support vector machine (SVM)-based algorithm for the automated detection of SWDs in the γ2R43Q mouse model of absence epilepsy. The algorithm first identifies putative SWD events using frequency- and amplitude-based peak detection. Four humans scored a set of 2500 putative events identified by the algorithm. Then, using predictors calculated from the wavelet transform of each event and the labels from human scoring, we trained a SVM to classify (SWD/nonSWD) and assign confidence scores to each event identified from 60 24-hour EEG records. We provide a detailed assessment of intra- and inter-rater scoring that demonstrates advantages of automated scoring.

**Results:** The algorithm scored SWDs along a continuum that is highly correlated with human confidence and that allows us to more effectively characterize ambiguous events. We demonstrate that events along our scoring continuum are temporally and proportionately correlated with abrupt changes in spectral power bands relevant to normal behavioral states including sleep.

**Significance:** While there are automated and semi-automated methods for the detection of SWDs in humans and rats, we contribute to the need for continued development of SWD detection in mice. Our results demonstrate the value of viewing detection of SWDs as a continuous classification problem to better understand ‘ground truth’ in SWD detection (i.e., the most reliable features agreed upon by humans that also correlate with objective physiological measures).

**Key Point Box:**

- Clinicians and researchers may benefit from an automated method of SWD detection that provides a framework for the quantitative description of SWDs and how they relate to other electrographic events.
- We present an algorithm for the automated, consistent, and rapid scoring of SWDs that assigns a confidence to detected events that is highly correlated with human scoring confidence.
- We characterize the human inter- and intra-rater consistency in the scoring of potential SWD events and compare them with the algorithm.
- Events along the scoring continuum generated by the algorithm are temporally and proportionately correlated with changes in spectral power bands relevant to behavioral states including sleep.

## Introduction

Spike-wave discharges (SWDs) are the signature electrographic (i.e. EEG) feature associated with absence epilepsy ^1,2^. These oscillations are thought to be associated with reverberations in thalamocortical networks and in rodents have a characteristic ∼6Hz oscillation ^3-5^. Identification of SWDs from EEG records is extremely important both in the clinic as a tool for the diagnosis of absence epilepsy ^6^ and in research. In the clinic, identifying the presence of seizure related events in a patient’s EEG may be sufficient to identify appropriate treatments. However, detailed scoring of epilepsy-related events may be required for better patient care ^7^ and to identify potential mechanisms of disease or to investigate the efficacy of drug treatments in research labs.

Manual scoring of SWDs is time-consuming and costly ^8^. In our experience, a skilled human may require 2-3 hours to score a 24-hour record. In both clinical and research settings there may be many hundreds of hours of EEG from humans or experimental animals that must be scored. As a result of the significant burden of scoring numerous long EEG records, research labs may only use snippets of EEG that are presumed to be representative of the full dataset. Selective cherry-picking of data could introduce unanticipated bias.

In addition to the time and cost associated with manual scoring of SWDs, inconsistencies between multiple scorers may be abundant ^3^. Such inconsistencies are not well characterized for SWD detection (we present a detailed example here), however inconsistencies have been described in the scoring of other nonconvulsive epileptiform events (e.g. interictal spikes) ^9^. Indeed, nonconvulsive epileptiform events are difficult to identify and classify for numerous reasons. First, the definition of ‘epileptiform’ is often vague ^10-12^ and is subject to disagreement. Clinicians and researchers typically use subjective criteria, such as that an event ‘stands out of the background’ ^13^, to force events into binary categories. Additionally, many electrographic features appear hidden in the time domain ^14^ and thus may be difficult to detect via visual inspection of EEG records. As we show, events can appear ambiguous to the same expert scorer such that, on repeated presentations, a scorer changes their mind about an event’s categorization. Such ambiguity is reminiscent of human perception of optical illusions such as the Necker Cube ^15^ and suggests that ‘ground truth’ (What do humans define as SWDs and how do those events relate to, or arise from, ongoing non-epileptic brain activity?) in identifying electrographic events may need to be considered in terms of confidence measures rather than certainty.

Thus, there is a need for the development of robust, automated methods for the detection of SWDs that allow for confidence-based scoring of events along a continuum that mirrors physiologically relevant EEG features and that matches human scoring characteristics. While algorithms have been developed for detection of SWDs ^16-19^, there has been little development of algorithms for the detection or quantification of SWDs in mice ^16^. This is an important gap because rodent models are currently the main research tools for understanding basic mechanisms of epilepsy.

We present a support vector machine (SVM)-based algorithm for the detection of SWDs in mice expressing a GABA_A_ receptor mutation (γ2R43Q) that causes absence epilepsy ^5,20^. This algorithm is much faster than a human, scoring a 24-hour record in ∼3 minutes. Importantly, this algorithm agrees well with human scoring and captures the statistical characteristics of human uncertainty. Additionally, and unlike human experts, it is 100% self-consistent. Finally, we show that events along the scoring continuum generated by the algorithm are temporally and proportionately correlated with changes in spectral power bands relevant to behavioral states including sleep.

## Methods

All use of animals in this manuscript conformed to the *Guide for the Care and Use of Laboratory Animals* ^21^ and was approved by the University of Wisconsin-Madison Institutional Animal Care and Use Committee.

### Animal colonies

Animals were bred from a colony maintained at the University of Wisconsin-Madison. In a paired breeding scheme, female wild-type (RR) mice of a C57BL/6J-OlaHsD background (Harlan, Madison, WI) were bred with male heterozygous mice expressing the γ2R43Q knock-in mutation in the same background. Heterozygous knock-in mice (RQ) experience SWDs concurrent with behavioral arrests ^5^ whereas homozygous mutants were almost never born. The γ2R43Q strain was provided by Dr. Steven Petrou (The University of Melbourne, Parkville, Australia).

### Computer Programing and Code Accessibility

All analyses, data processing, and event classification was done in Matlab (Mathworks, Natick, MA). The final version of this algorithm was executed with Matlab 2017b and code is available at the GitHub repository: https://github.com/jessePfammatter/detectSWDs.

### EEG implantations, recording, and data

EEG electrodes were implanted as described in ^22^. Briefly, male heterozygous knock-in (RQ) and wild-type (RR) mice were implanted at p24 under isoflurane anesthesia (1-2% in 100% O2). Each animal was implanted with gold plated miniature screw electrodes over the right and left frontal and parietal cortices, and one over the cerebellum as reference. Two vinyl-coated braided stainless-steel wire electrodes were placed in the nuchal muscle for electromyogram (EMG) recording. The EEG and EMG leads were wired to a head cap, which was affixed to the skull with dental acrylic.

After recovery from surgery (2-3 days), animals were connected to a multichannel neurophysiology recording system (Tucker-Davis Technologies, TDT, Alachua, FL, USA) to continually sample EEG and EMG signals at 256 Hz (digitally bandpass filtered between 0.1 and 100 Hz) for up to fourteen days. Offline, data were notch filtered at 60 Hz using a Chebyshev Type II digital filter and high-pass (>2 Hz) filtered with a Chebyshev Type I digital filter. We selected these infinite impulse response (IIR) filters because of their highly specific frequency attenuation without ripple in unintended frequencies, acceptable phase shift given our largely time-frequency based algorithm, and computational efficiency (see Supplemental Figure S1 for frequency and phase responses of these filters). EEG signals were then normalized using the following variation of a z-score normalization: First, we fit a Gaussian distribution to the all-points histogram (calculated using the histfit() function with bins equal to the floor of the square-root of the number of data points in each EEG signal) of each 24-hour record using least-squares minimization via Nelder-Mead Simplex optimization ^23^ as implemented by the fminsearch() function in Matlab using the mean and standard deviation of the 24-hour EEG as starting guessing for the optimization. We then normalized the 24-hour record by subtracting the mean from each data point and dividing by the standard deviation of the model fit. The resultant normalization is different from the standard z-score normalization in that the mean and standard deviation are calculated from estimating the Gaussian portion of the signal via the best-fit Gaussian model (using sum of squared errors) rather than empirically calculating the mean and standard deviation. We employed this normalization technique in order to comparatively normalize the EEG from healthy and epileptic animals alike. High amplitude EEG signal such as convulsive seizures or high amplitude spikes produce EEG data with non-Gaussian all-points histograms and thus poor results from the Maximum Likelihood Estimates of the mean and standard deviation. Additionally, more simplistic methods such as a median normalization did not perform as well and fundamentally do not attempt to normalize the data to the ‘normal’ component of the EEG signal in the same way as our method. All analyses used a frontal EEG channel.

### Automated Event Detection

We used a two-stage algorithm for the detection and classification of SWDs. In Stage 1, we automatically detected putative SWDs using a frequency and amplitude threshold-based approach. The parameters we selected for this first stage were set heuristically and intended to be ‘all inclusive’ of potential SWD events at the cost of also finding false positives, which were filtered out in Stage 2. First, we detected all peaks that were a) above 3 standard deviations from the mean of the normalized EEG signal and b) that had a peak greater than 3 standard deviations from the mean of the derivative signal (the negative of the first derivative of the normalized signal calculated with the gradient() function and normalized again with the same procedure as the original signal) that preceded (within 60 ms) the peaks identified in the normalized EEG. Then, sets of peaks were grouped into ‘possible SWD events’ if their frequency was between 3-11 Hz. This procedure selected events in order to minimize the number of false negatives as compared with unaided human scoring (i.e., no information from Stage 1) while also remaining within previously published guidelines for the characteristics of SWDs in mice ^24, 5^ and rats ^4, 3^.

### Manual Event Classification

Completely unaided from our computer algorithms, scorers (S1 and S4: RKM and JAP) each manually scored five 24-hour EEG records (n = 4 RQ, n = 1 RR) for the presence of SWDs. Scorers marked epochs (4s) containing SWDs using Svarog v1.0.10 (Braintech, Ltd., United Kingdom). We use these manual scores to help validate the performance of both our Stage 1 and Stage 2 algorithms.

We also presented 2500 putative SWD events identified by the Stage 1 detection method to each of four human scorers (S1-S4: RKM, EPW, MVJ, and JAP) including an epileptologist with over 20 years of experience (RKM) for manual classification. Of these, 2050 were randomly selected unique events from 10 24-hour EEG records from five animals (3 RQ and 2 RR). From these we selected 50 events to be repeated 10 times each. Each of the resultant 2500 events were shown to the human scorers in random order. Each event was presented as an 8-second length of normalized single-channel EEG with the putative event highlighted in yellow. Scorers classified events as either SWD or nonSWD and were given as much time as they needed to view/review/reclassify events. All human scorers were skilled in the identification of SWDs but were blind to the genotypes, and two were blind to the presence of repeated events. None of the scorers were aware of which events were repeated because the order of events was random.

EEG/EMG signals were also manually scored for sleep stages (wake, NREM, or REM) (see Fig. 3) in epochs (4s) with Serenia Sleep Pro software (Pinnacle Technology, Lawrence, KS, USA) following criteria similar to Nelson et al, (2013)^22^. Stretches ≥ 8 seconds (2 epochs) were required for valid transitions.

### Event Predictor Variables

From the putative events detected by Stage 1, we extracted 12 predictor variables for use in Stage 2 of the automated algorithm: event classification with a Support Vector Machine (SVM). These variables were calculated from four scale ranges from the wavelet transform (using the cwt() function specifiying ‘amor’ for the Morlet wavelet) of each event. These ranges were chosen to target the characteristic ∼6Hz frequency of SWDs in mice and several of its harmonic components ^3-5^. For each of the four frequency bands (4.4-8.2 Hz, 8.8-16.4 Hz, 17.6-32.8 Hz, and 35.1-65.5 Hz) we summed the absolute value in each bin across the duration of each event (9 bins per band) and calculated the mean, standard deviation, and maximum values for each event (four frequency bands * three statistics = 12 predictors).

### Automated Event Classification

For Stage 2 of the algorithm we developed an SVM-based automated method to classify SWD events identified in Stage 1 using the 12 predictors. In brief, a SVM is a machine learning-based classification method that aims to separate groups of events after being trained with set of ‘known’ labels. The boundaries of the classification space are defined by events assigned as support vectors. The location of each event relative to the nearest support vectors can be used for classification of events and the distance from each event to its nearest support vector can be used as a proxy for confidence in the assigned classification of each events. Readers unfamiliar with SVM may find a suitable introduction in ^25^ or ^24^. We trained the SVM (fitcsvm() function) using the labels provided by the human scorers (10000 labels, 2500 per scorer for 2050 unique events) where the algorithm created a weighted response for each unique event based on the average human label across all four scorers. In our preliminary testing of the algorithm and upon visual inspection of the predictor variables for these events, we choose a Gaussian kernel for the SVM ^24^. In order to choose the proper hyperparameters for the SVM (kernel scale, box constraint, and cost matrix used in fitcsvm()), we employed a grid-search based optimization process. We trained the SVM under each unique combination of the following parameters; kernel scale = [1, 5, 10, 15, 20], box constraint = [5, 10, 15, 20, 25], and a cost matrix of [0, 1; x, 0] where x = [1, 1.5, 2, 2.5, 3]. We then chose hyperparameters with the following guidelines: A) We aimed to minimize the standard deviation of the agreement among the human scorers and the SVM in order to minimize individual scorer bias. We chose this method as opposed to maximizing the average agreement between the human scorers and the SVM because that quantity can be maximized to a significantly higher value than the agreement within human scorers, resulting in model overfitting and poor performance on novel data. B) To avoid overfitting, we calculated the 5-fold cross validation, which partitions the data into multiple subsets and tests model fit on each subset ^24^, between the SVM and the training labels. Because Matlab uses a stochastic fitting process for the generation of the SVM, we repeated SVM fitting and calculation of the 5-fold cross validation 10 times and took the standard deviation of 5-fold cross validations ^26^. Minimizing this quantity allowed us to select a model that performs similarly on numerous randomly chosen subsets of data. C) The model should match human uncertainty in order to generate accurate confidence-based model scores (distance from each event to the nearest support vector). We aimed to maximize the R^2^ between the model scores and the weighted human responses for each of the 50 repeated events (not the entire 2500 events) within the data set (i.e. intra-rater reliability) which produces a model with high fit to the human uncertainty (the 50 repeated events are our predictors of human uncertainty in SWD scoring). While these parameters are important, their optimal set point is unclear given that variability between human scorers suggests that ground truth labels may not exist for these data. Here, we used the following hyperparameters: kernel scale = 10, box constraint = 10, cost matrix x = **1.5**.

In order to better understand the relationship of the 12 predictor variables calculated from the wavelet transform of each event to the classification scores produced by the SVM, we performed a LASSO regression ^27^. While this analysis does not allow direct understanding of the classification space of the SVM, such direct understanding is difficult to achieve and therefore methods that allow some intuition as to the relative importance of predictor variables in the system are useful. We allowed the LASSO regression 100 iterations with cross validated (10-fold) fits and selected the final coefficients from the model with a lambda value (indicating lowest cross validation error) that was one standard deviation larger than the model with the lowest lambda value.

In order to highlight the differences between the human and SVM scoring of the 2500 events identified by Stage 1, we calculated receiver operating characteristic and precision-recall curves between each scorer and the algorithm, and a table showing pairwise agreement between the human scorers, the SVM trained with K-means ^28^ labels (generated using the kmeans() function from the 12 predictor variables for each event), the final SVM, and the human consensus labels. In addition, we calculated the sensitivity, specificity, and precision between the human consensus labels and the final SVM (for the 2500 events identified by Stage 1). In order to demonstrate the performance of the algorithm in comparison to human scoring (2 scorers) of an unannotated (no information from Stage 1) and out-of-training data set (five, 24-hour EEG recordings), we calculated confusion matrices from the epoch-by-epoch scoring of SWDs identified by humans (S1 and S2) and the Stage 1 events, and SWDs identified by humans (S1 and S2) and the Stage 2 (final SVM) scores. From these confusion matrices we calculated the 12 event predictor variable values for each event within three categories; Stage 2 true positives (TP, events within epochs classified by Stage 2 the humans as SWDs), Stage 1 false negatives (FN, events within epochs classified by humans as SWDs but missed by Stage 1), and Stage 2 true negatives (TN, events within epochs classified by Stage 2 and the humans as nonSWDs). We then calculated a multivariate analysis of variance (MANOVA, using the manova1() function) to demonstrate if the means of the 12 predictor variables collectively varied between the three test groups (Stage 2 TP, Stage 1 FN, and Stage 2 FN).

### Application of the algorithm and preliminary testing of the SWD output scores

We applied the algorithm to classify SWDs in 60 24-hour EEG records from 8 RQ and 3 RR animals including 45 records that were not used for algorithm development (i.e., out-of-sample data). Using the output of the algorithm we calculated the event-triggered average: (i.e., the average of all segments of one signal aligned to the time of events in another signal) of 50 random examples of strong SWDs (distance to support vector of >1 to 2), moderate SWDs (>0.5 to 1), weak SWDs (>0 to 0.5), weak nonSWDs (<0 to −0.5), moderate nonSWDs (<−0.5 to −1), and strong nonSWDs (<−1 to −2), and 50 random locations not identified by our algorithm to delta (0.5-4 Hz), theta (6-9 Hz), sigma (10-14 Hz), and gamma (25-100 Hz) power surrounding each event during lights-on and lights-off periods.

## Results

We found that events identified by our algorithm as SWDs in γ2R43Q animals have a characteristically strong and continuous power in the ∼6 Hz frequency range during the entire discharge event, (Fig. 1) consistent with previous descriptions of SWDs in these animals ^5^. Additionally, these events have a reliable ‘fingerprint’ in the time-frequency decomposition (Fig. 1).

**Figure 1:**
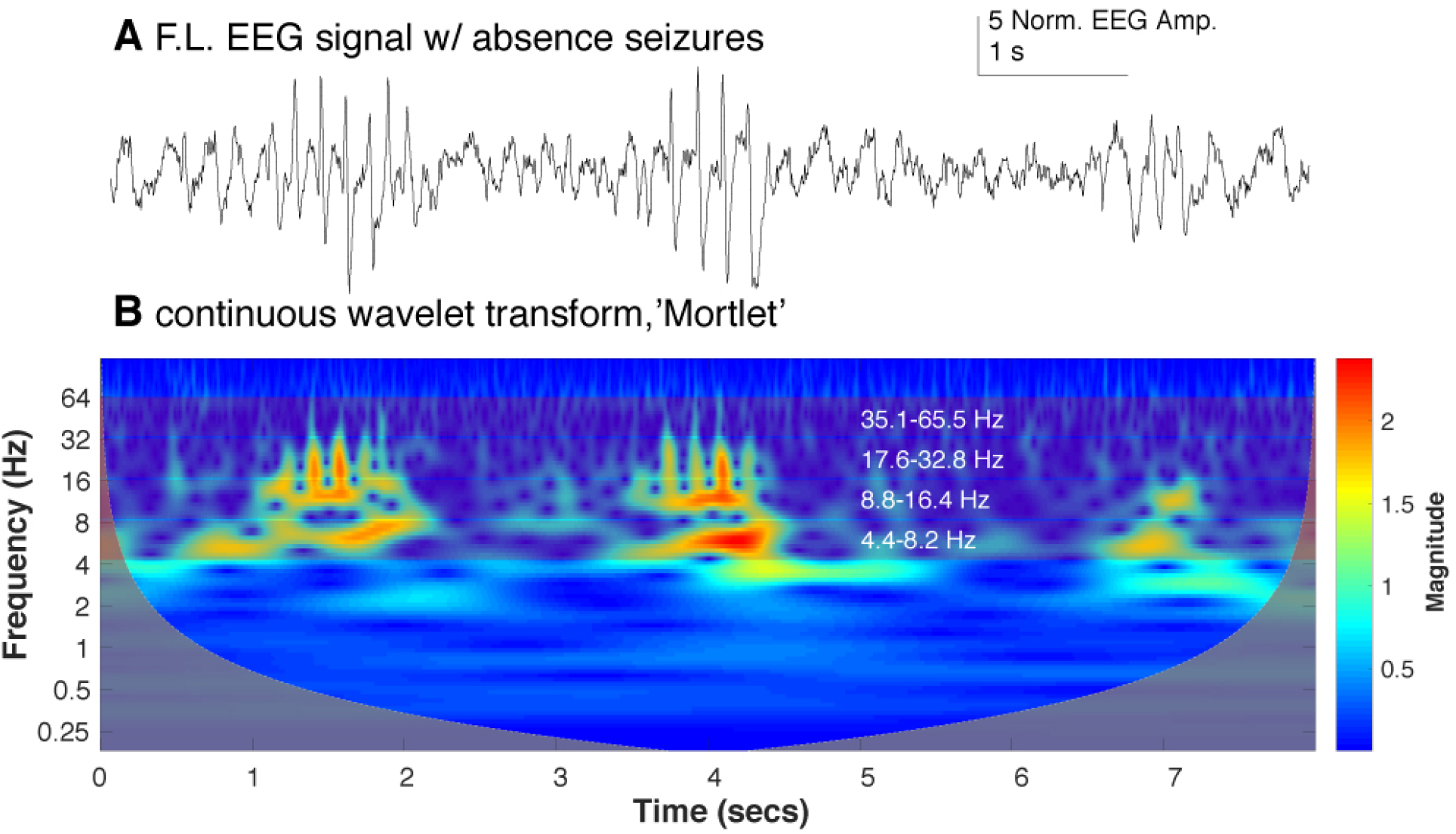
**(A)** An EEG trace from the frontal left channel of a mutant γ2R43Q mouse showing three putative SWD events detected by Phase 1. **(B)** The wavelet transform scalogram of the trace in A. We use the maximum, average and standard deviation of the scalogram amplitude from four bands (∼4.4-8.2 Hz, ∼8.8-16.4 Hz, ∼17.6-32.8 Hz, and, ∼35.1-65.5 Hz shown as bands with white frequency labels) to generate 12 predictors to train the SVM-based classification algorithm.

### Manual Scoring and Inter- and Intra-Rater Reliability

Overall, human scorers were quite variable in their scoring. Out of the 2050 unique events, the four human scorers were in complete agreement (i.e., consensus labels) for 57% of the events (1159 events, 940 nonSWD and 219 SWD). Figure 2 shows a characterization of human scoring of the 2500 events presented as 2D cross-sections in predictor space for 6 of the predictor variables used in the SVM. Further characterization of the inter- and intra-rater reliability are shown in Figure 3, which demonstrates the difference in rating trends of the human scorers. Figure 3B shows 8 example events pulled from the 50 events presented to each human scorer 10 times. For each of the 2500 events presented to human scorers, agreed-upon SWD events have high values for both the maximum values in 4.4-8.2 Hz and 17.6-32.8 Hz ranges whereas agreed-upon nonSWD events have a low maximum value for these bands (Fig. 3B). Intra- and inter-rater reliability data are shown in Figure 3C-E. Inter-rater reliability on the 50 repeated events (RE) data (Figure 3D) agreed with the trend in distribution of human agreement for all of the 2500 events presented (low-resolution data). Average intra-rater reliability across all scorers was 100% for 13 of 50 repeated events (Figure 3E). Thirteen of the 50 RE had an average intra-rater agreement of 90% or less across all scorers. For individual scorers, intra-rater agreement was as low as 50% for the most ambiguous events. Average Intra-rater reliability was 94% across all scorers and all 50 repeated events.

**Figure 2:**
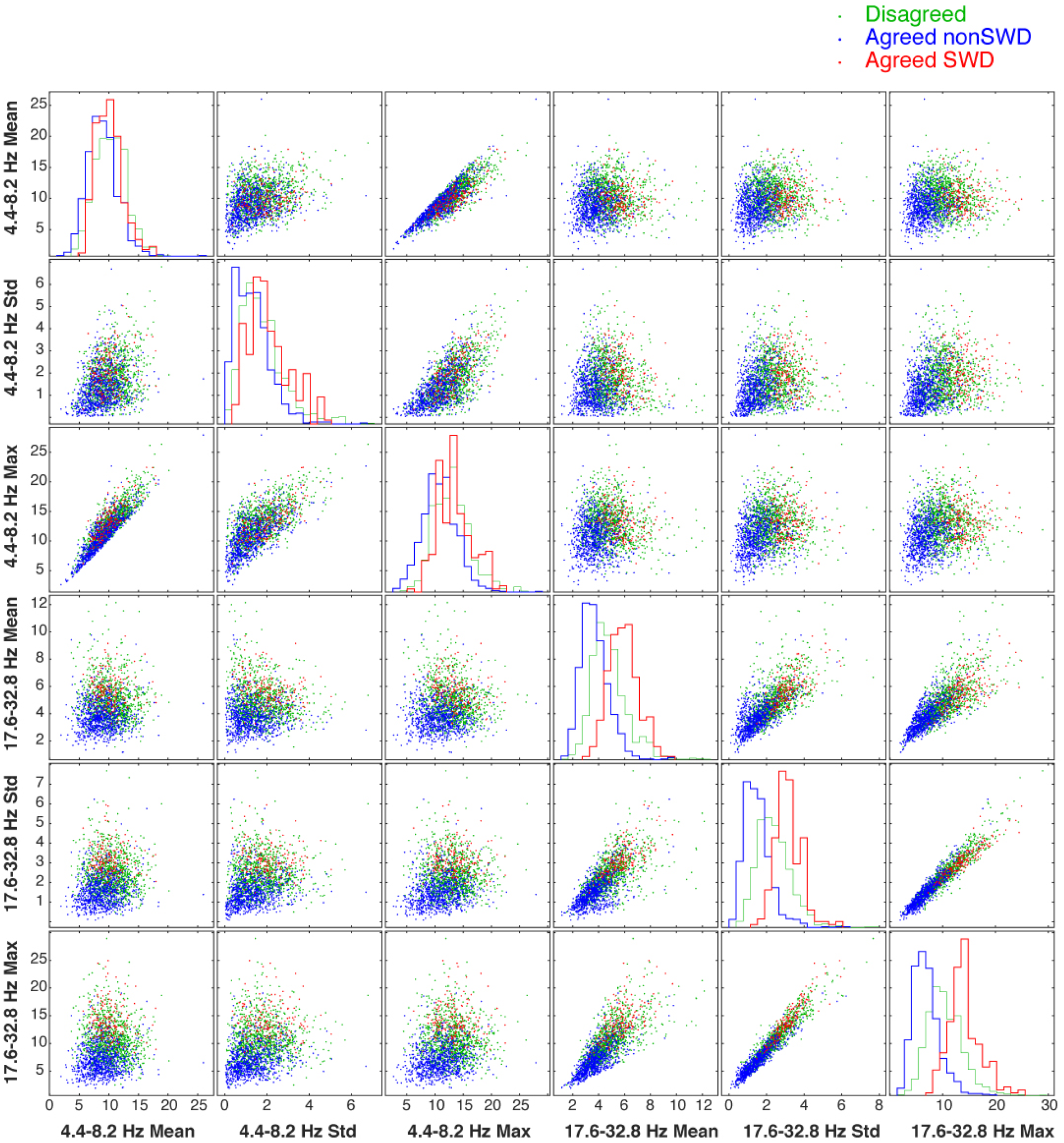
Each of the 2500 human-scored events labeled by agreed SWD (red), consensus agreed nonSWD (blue), and consensus disagreed (green) shown as a 2D cross-sections in predictor space for 6 of the 12 predictor variables used to characterize a SWD event. The diagonal shows the histograms for each category of label for each of the 6 predictor variables. Notice how disagreed-upon events tend to align with SWD rather than nonSWD events.

**Figure 3:**
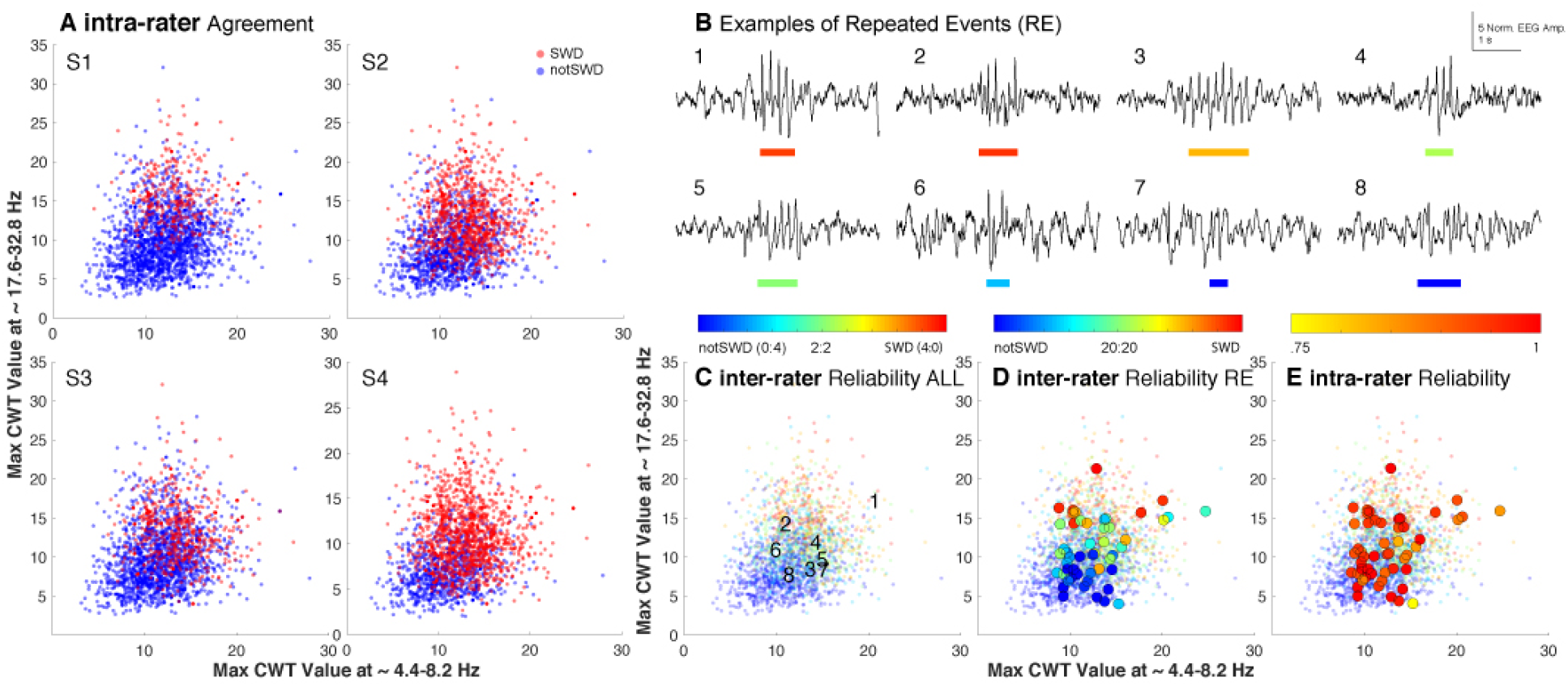
A characterization of the variability of human classification of SWD events. **(A)** Each of the 2500 events as labeled by human scorers 1-3. Note the dramatic variability in labeling between each of the scorers. **(B)** examples of eight events identified as ‘putative’ SWD events by the first stage of the algorithm. The colored bar below each trace represents the start and end of each event, where color shows the degree of agreement between human scorers. This color scale matches that above Panel C. **(C)** Each of the 2500 putative SWD events presented to each human scorer, plotted by the maximum value of the wavelet transform from ∼ 4.4-8.2 Hz versus the maximum value of the wavelet transform from ∼ 17.6-32.8 Hz. The numbers mark locations the 8 example events from panel B. **(D)** The average ***inter-rater*** agreement for each of the 50 repeated events (RE, large circles) plotted over the same data. Note that the agreement between raters approaches a coin flip (green) near the intersection of the strong SWD and nonSWD events. **(E)** The average ***intra-rater*** reliability for the same 50 repeated events as shown in panel D. Note the shifting scale associated with the color bars above panels C, D, and E.

### Comparing Automated and Manual Scoring

While only one of many possible clustering methods, K-means, a more traditional clustering method, failed to partition the data similar to the human scorers (notice the ‘fuzziness’ in SWD/nonSWD scoring in Figure 3A) but rather drew a hard line between SWD/nonSWD events (Fig. 4A). The output of the SVM trained with human labels demonstrates a similar ‘fuzziness’ in scoring as human scorers (Fig. 4B). In fact, the SVM classification associated with events (e.g. distance to the nearest support vector) is strongly correlated with human inter-rater reliability (Fig. 4C). In comparing labels from our final SVM algorithm and the human consensus labels, the sensitivity (equal to the number of true positives, the SWD events agreed upon between SVM and the human consensus labels, divided by the total number of SWD events from the human consensus labels) was 0.98. The specificity (equal to the number of true negatives, the nonSWD events agreed upon between SVM and the human consensus labels, divided by the total number of nonSWD events from the human consensus labels) was 0.95. The precision (equal to the true positives divided by the sum of the true positives and false positives, the number of events classified as SWD by the SVM but nonSWD by the human consensus labels) was 0.83. The values for sensitivity, specificity, and precision presented here were calculated using events identified by Stage 1 of the algorithm and while these are useful metrics they should not be interpreted as the sensitivity and specificity of the algorithm as compared to unaided human scoring of SWDs. Figure 4D shows the coefficients for the LASSO regression. The absolute value of these coefficients indicates the relative importance of each variable whereas the sign indicates whether a large versus small value of that variable was most predictive of the SVM classification scores. Negative coefficients do not indicate that the magnitude for those variables were lower for SWDs than nonSWDs on average. Figure 4E-F shows the receiver operating characteristics curves and precision recall curves between the 2500 events scored by each of the human scorers and the SVM scores. The average agreement between humans was similar to that between humans and K-means but lower than the average agreement between the SVM and the humans. The standard deviation of the agreement between the SVM and the human scorers is lower than between the human scorers themselves indicating that the SVM agrees with the better than the humans agree with themselves (Fig. 4G). Figure 4H shows a comparison between performance of the scoring of five unannotated (without computer assistance) 24-hour EEG records by humans (S1 and S4) and the SVM. The first stage of our algorithm captured a large number of the epochs marked as containing and SWD by the humans (1933 of 1962 epochs). Of those epochs containing SWD events identified by humans but entirely by Stage 1 of the algorithm (Stage 1 FN, 290 collective epochs containing 244 unique events), events within those epochs have significantly different characteristics (values for predictor variables used in the SVM) than those events within epochs identified as Stage 2 TP (1490 collective epochs containing 1150 unique events) or Stage 2 TN (240227 collective epochs containing 3143 unique events) (MANOVA dimension = 2, *P-values* < 0.001, Figure 4H).

**Figure 4:**
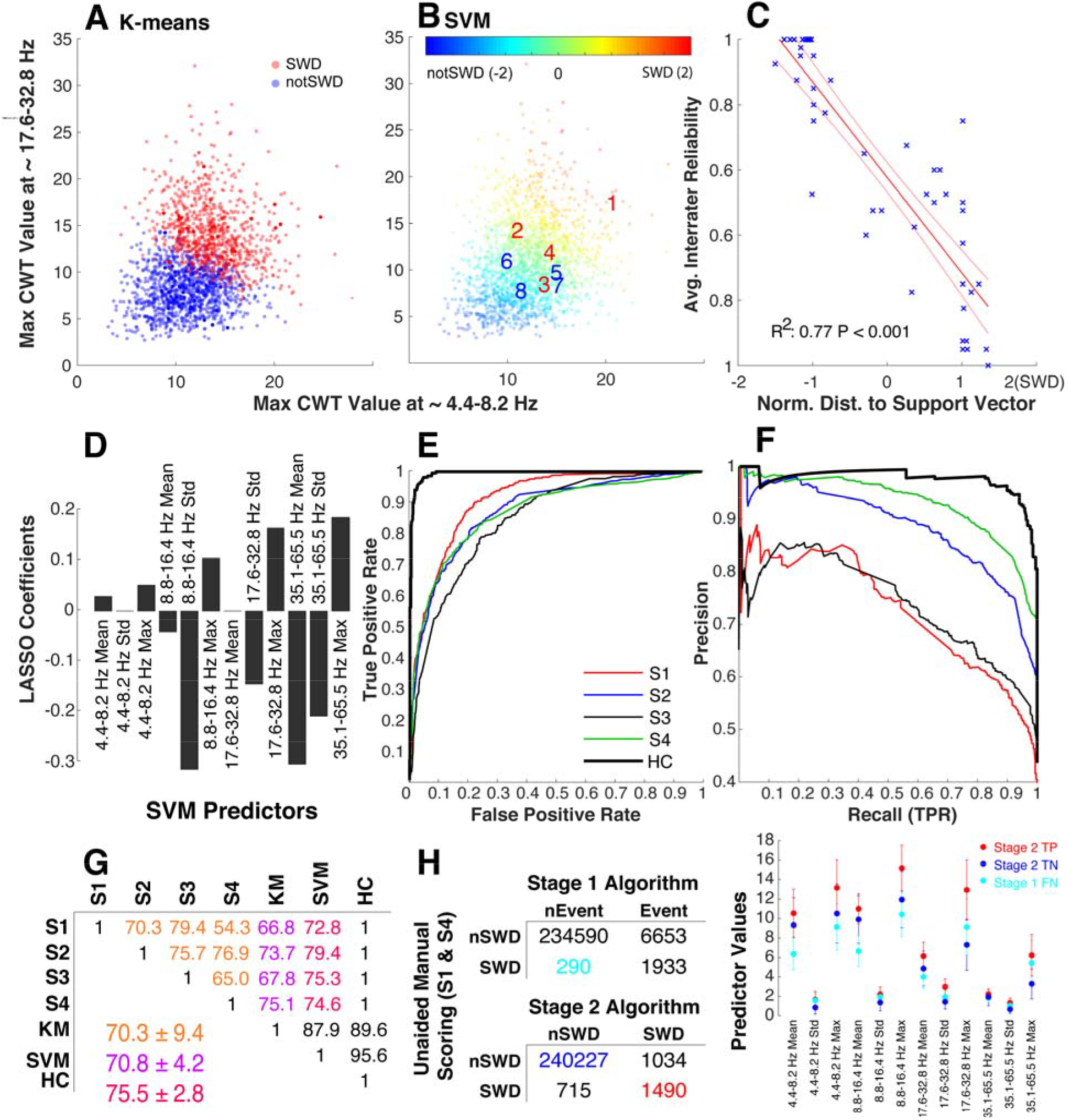
SVM-based approach to SWD classification. **(A-B)** each of the 2500 potential SWD events displayed by the maximum value of the wavelet transform from ∼ 4.4-8.2 Hz versus the maximum value of the wavelet transform from ∼ 17.6-32.8 Hz. **(A)** shows events as clustered by K-means clustering using the 12 wavelet transform predictor variables. **(B)** Events labeled by a SVM trained with the human scoring labels and fit with a Gaussian kernel. The numbers 1-8 are overlaid to show the SVM’s classification of the 8 example events from Figure 2. **(C)** The relationship between the distance to nearest support vector (x axis, 2 is more likely SWD) and the average intra-rater reliability (from Fig. 2C) for the 50 repeated events. **(D)** The coefficients from a LASSO regression which highlights the relative importance (absolute value of coefficients) and relative magnitude (sign of coefficients) of the variables as compared to the SVM scores. **(E)** Receiver operating characteristic and **(F)** precision-recall curves showing the performance of each human scorer and the human consensus labels (HC) as compared to the SVM scores. **(G)** The agreement matrix for classification of 2500 potential SWD events as classified by four human scorers (S1-S4), an SVM trained with the KM labels, and the human consensus data (HC, the events on which all human scorers agreed). Note that the agreement between all of the humans (orange) was comparable to that between the SVM classification and the humans (red) although the agreement between the SVM and the humans shows much lower variability than the agreement between the humans alone. **(H)** Confusion matrices comparing the epoch-by-epoch scoring of events identified by Stage 1 of the algorithm (top matrix) and events classified as SWD by the Stage 2 SVM (bottom matrix) and S1/S4 scoring of five 24-hour EEG records representing two RQ and two WT animals. The Youden’s J statistic (sensitivity + specificity −1) between S1 and S4 equals 0.80. S1 and S4 scored these files without the any information from the algorithm regarding putative SWD events. Events within epochs identified as Stage 2 TP (red), Stage 1 FN (cyan), and Stage 2 TN (blue) have significantly different multivariate means for the 12 predictor variables used by the SVM for event classification (lower-right plot).

We present graphical results for the optimization of our tunable SVM parameters in the Supplemental Figure S2.

### Application of the Algorithm

We applied the algorithm to analyze SWDs in a set of 60 24-hour EEG recordings from RR and RQ animals, 50 of which were not used for SVM development. The algorithm analyzed these records in ∼3 minutes per 24-hour record. Figure 5 shows the application of our algorithm on a representative 24-hour record from a RQ animal. Additionally, using the output of the algorithm on an RQ animal, we demonstrated the relatedness of SWD and nonSWD events (as labeled by the SVM) to sleep staging. We found strong and abrupt relationships between SWDs and each power band that sometimes preceded the SWD by several minutes (Fig. 5D-K).

**Figure 5:**
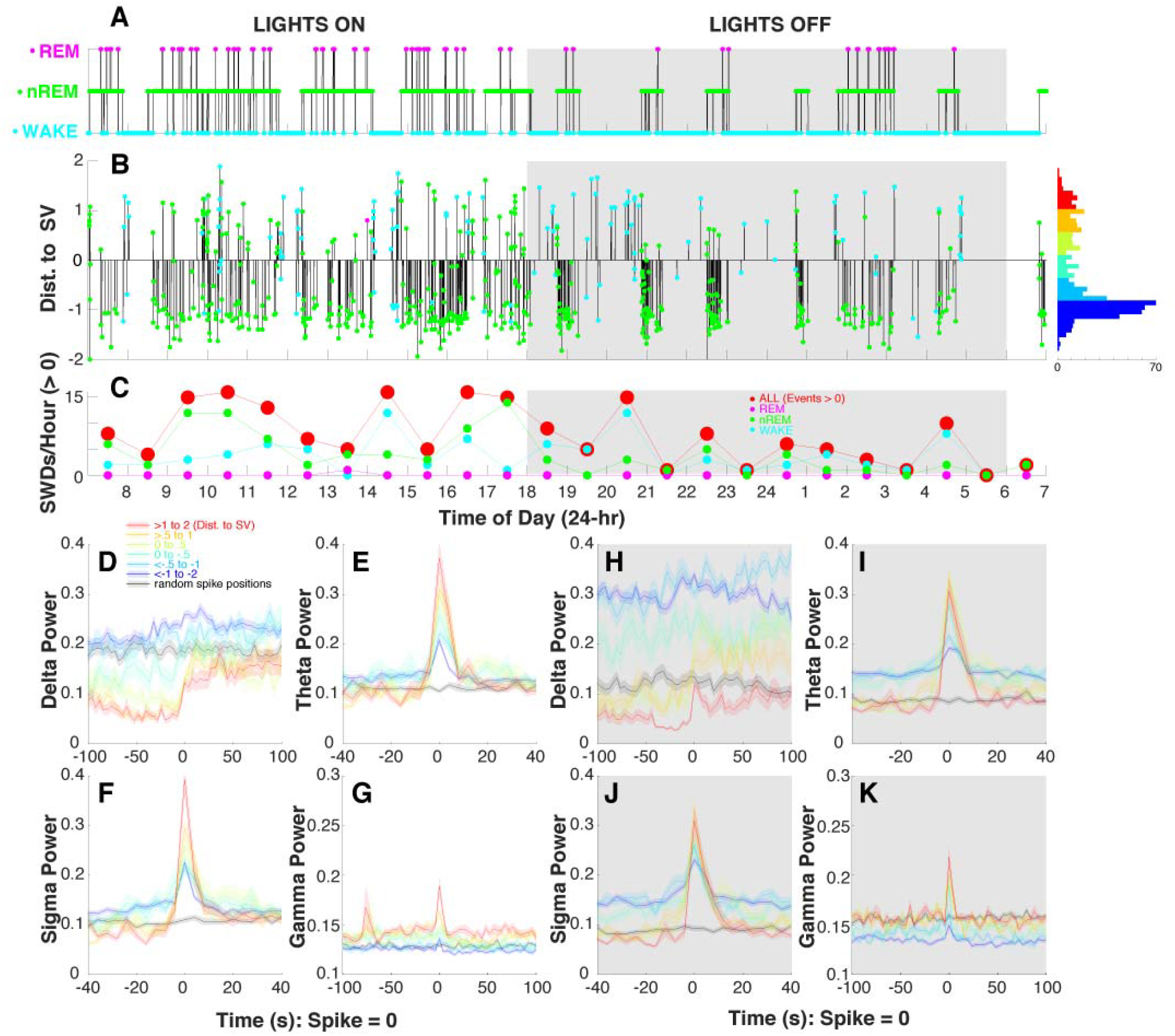
Implementation of the SWD detection algorithm on a 24-hour EEG recording from a mutant γ2R43Q mouse with absence epilepsy and a demonstration of the high-resolution data provided the ‘distance to SV’ measure associated with each identified event. **(A)** A hypnogram showing WAKE (blue), nREM (green) and REM (magenta) epochs (4 s) manually scored by RM. **(B)** A visual representation of the events identified by the SVM-based algorithm. The vertical height of each bar represents the distance to nearest support vector where values > 0 are likely to be SWDs whereas events < 0 are likely nonSWD events. The color at the end of each vertical bar shows the sleep stage associated with the identified event. The histogram on the right of panel C shows the distribution of event scores and colors are representative of the categories of events used in panels D-K. **(C)** shows the average number of SWDs (score > 0) per hour for all sleep stages (red, larger circles), WAKE, nREM, and REM sleep stages. Notice the pattern of increased SWD events during lights on periods (07:00 - 18:00, normal sleep time for mice) as compared to lights off. The event-triggered average of delta **(D, H)**, theta **(E, I)**, sigma **(F, J)**, and γ **(G, K)** power (calculated in 4 s epochs) for events at several sets of distances from the nearest support vector where redder colors are more likely to be SWDs (red: >1 to 2, yellow: >0.5 to 1, lime: 0 to 0.5, green: 0 to −0.5, cyan: <−0.5 to −1, blue: <−1 to −2). Note that the color scale in this figure matches the color scale in panel B of Figure 3. The black trace in each of these panels shows the null-hypothesis: the event-triggered average of each power value for 100 randomly selected epochs in the EEG record. Events with the largest positive distance to SV (red) show a strong relationship to all power bands during lights on (white panels) and lights off (grey panels) and this relationship deteriorates as event distances decrease. Note that all events identified by the algorithm show some sort of temporal relationship with theta and sigma power as compared to randomly selected EEG locations. Further investigation is needed to identify the relationship of each group of these events to absence epilepsy.

We found an average of 737.51 ± 482.28 events per 24-hour RQ record (n = 8 animals, 45 24-hour records), of which 194.09 ± 169.37 of were classified as SWDs (> 0 SVM score). In RR animals (n = 3 animals, 14 24-hour records), we found 409.64 ± 207.28 possible events per 24-hour record, of which 26.79 ± 21.18 were classified as SWD events. The events identified as SWDs had a significantly lower SVM score in RR animals (0.797 ± 0.445) than in RQ animals (0.519 ± 0.376, t = 24.39, df = 17410, *P-value* < 0.001). Despite having a lower average SVM score, SWDs identified in RR animals show strong visual SWD characteristics (Fig. 6A) and show similar temporal relationships with delta, theta, sigma, and gamma power bands as events from RQ animals (data not shown). Despite being very similar, events from RR animals do have a significantly different multivariate profile for the 12 cwt-based predictor variables than the RQ animals (MANOVA df_Chi-squared_ = 12, Chi-squared value = 984.27, *P-value* < 0.001, Figure 6B). However, these differences appear to be relatively small compared to those differences observed between TPs and TN/FNs presented in Figure 4H.

**Figure 6:**
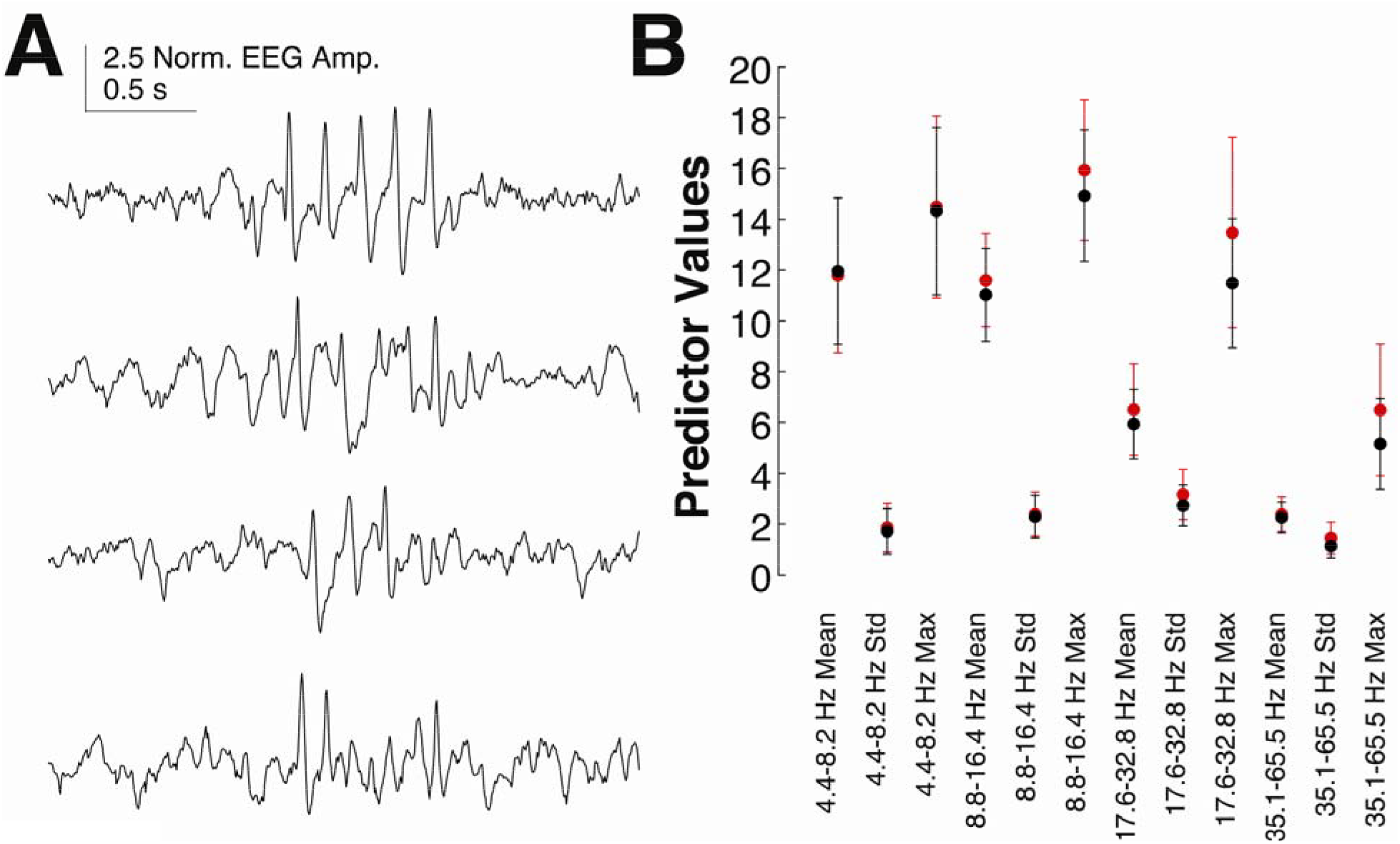
Four SWD-like events from a 24-hour record of a wild type (RR) animal as identified by the SWD detection algorithm. **(B)** Means and standard deviations for the 12 predictor variables calculated for SWD events found by our algorithm in RR (black circles, n = 3 animals, 14 24-hour records) and RQ (red circles, n = 8 animals, 45 24-hour records) animals.

## Discussion

We have developed an automated, SVM-based algorithm for the detection of SWDs that leverages human agreement in the scoring of machine-preselected events to develop a continuous, confidence-based score in addition to a binary classification (SWD/nonSWD). The algorithm was trained using a dataset with 2500 putative events and labels (SWD or nonSWD) from four expert human scorers, using 5-fold internal cross validation, and tested against the performance of two human scorers (S1 and S4) on multiple, unannotated (no computer information), and out out-of-training data records. The resultant SVM agreed with humans as well as they agreed with each other when scoring computer pre-selected events (∼70%) but, unlike the other classification technique that we investigated (K-means, Fig. 4A), the SVM did not draw sharp boundaries in predictor space between regions containing SWDs vs nonSWDs. Instead, it drew fuzzy, tightly interleaved boundaries similar to those of human scorers. The algorithm also correlated well with the uncertainty of individual humans when presented with repetitions of the same waveforms, thus capturing ∼77% of the ambiguity in human perception of SWD waveforms (Fig. 4C). Although other groups have developed algorithms for the detection of SWDs ^16-19^, to the best of our knowledge, no other group has designed an algorithm to mirror human confidence in scoring. Indeed, we focus our most significant attention on this component of the algorithm as we feel it is perhaps the most important contribution of this manuscript to the current state of event detection in absence epilepsy.

Although the method works quite well on the γ2R43Q model of absence epilepsy, we have yet to fully test it in other models of absence epilepsy. Continued development will include testing and expansion to other animal models as well as to human EEG. We anticipate that Stage 1 (event detection) should work quite well for most types of absence seizures exhibiting a ∼6 Hz character (as occurs in rodents). The frequency range of interest can be easily adjusted for different data sets. However, for Stage 2 (the SVM classification), some retraining of the algorithm may be necessary, using human scoring on a very small subset of data that are representative of the different types of subjects/treatments being analyzed. In this current version, we selected a set of optimization parameters and SVM kernel that best fit the characteristics of human scoring. In general, modification of the hyperparameters or kernel can make a SVM more generalizable ^25^. Continued development of this algorithm may also include components shown to be important for other SWD detection algorithms such as using phase information from the wavelet transform such as was done in ^16^. Additionally, some concern may exist about the nonstandard normalization procedure used in this algorithm. Although we are confident in this normalization, and are not removing or cleaning the data through this procedure (besides the stated filters), the matter certainly deserves more attention and is one focus of our ongoing research.

Despite its limitations, we have used this algorithm to highlight a few important aspects of SWDs that would have been difficult to identify solely with human classification. Our data show that events along the spectrum of the SVM score (and thus SWD confidence) show temporal and proportional correlations with abrupt changes in EEG power bands. While these correlations are most prominent in SWDs with the highest SVM scores, events with lower scores (and even those below threshold) continue to show correlations, albeit with smaller effects. Qualitatively similar temporal patterns were observed in all of the R43Q animals analyzed in this study (data not shown). Thus, the SVM reveals physiologically-relevant changes in brain activity that are commonly used in the scoring of sleep and shows that these changes are temporally linked to, and could potentially be used to predict, the occurrence of SWDs. Importantly, our findings align with previous work showing shifts in phase-amplitude coupling up to a minute prior to SWDs ^29^, and that SWDs are related to sleep timing ^30^. Additionally, our algorithm includes SWDs and SWD-like activity of short duration which are discarded by most studies ^16^. We also identified SWD-like events in wild-type mice, consistent with the hypothesis that SWD events represent a corruption of otherwise normal brain processes ^31^. Given this hypothesis, finding examples of SWDs in wild-type mice with low intensity (i.e. smaller distance to nearest SV) is to be expected. Additionally, ^32^ found that 8 of 27 strains of wild type mice had SWDs including several from the C57-related backgrounds. While low intensity events show temporal relationships with several EEG power bands that coincide with confident SWD events (data not shown), further research is needed to understand if the SWDs observed in RR animals are related to the same processes as those that generate SWDs in absence epilepsy or if they are representative of other normal physiological processes.

## Acknowledgements

We would like to thank Kile P. Mangan for scoring SWDs, Eli Wallace for scoring sleep, and Rebecca Willet for helpful conversations regarding SVMs. The data analyzed were collected by KPM and Aaron Nelson in the lab of Chiara Cirelli. Antoine Madar and Eli Wallace provided invaluable feedback during the development of this work. Rachel Bergstrom and anonymous reviewers provided valuable reviews which improved this manuscript. This work was funded in part by grants from the National Institute for Health (RO1 NS075366, MVJ) and the Department of Defense (PR161864, RKM).

## Conflicts of Interest

The authors report no conflicts of interest.

## Ethical Publication Statement

We confirm that we have read the Journal’s position on issues involved in ethical publication and affirm that this report is consistent with those guidelines.

---

**Supplemental Figure S1:**
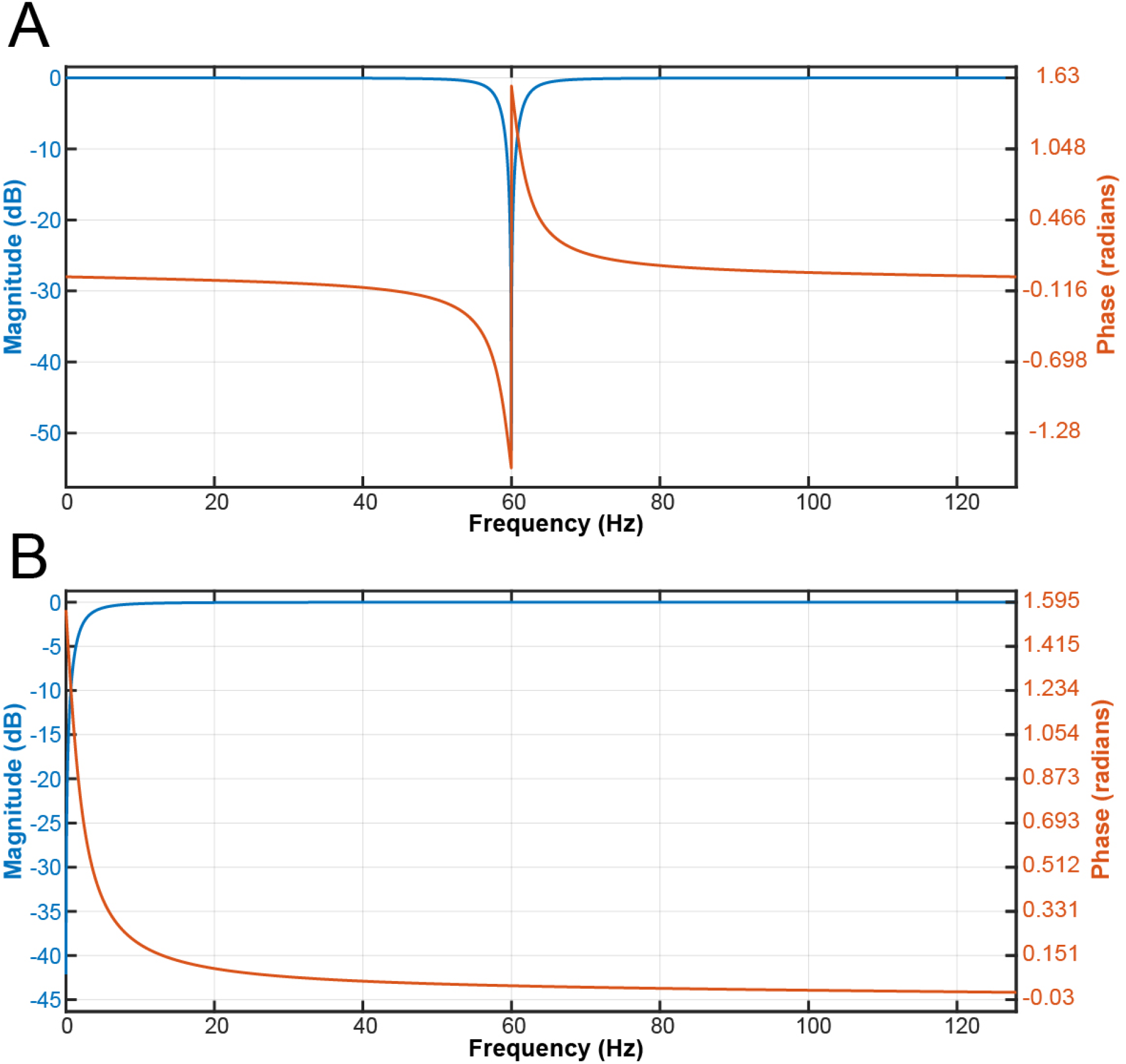
Amplitude (blue) and Phase (orange) response of the 60 Hz notch (Chebyshev Type II) and high-pass Chebyshev Type I) filters used in the algorithm.

**Supplemental Figure S2:**
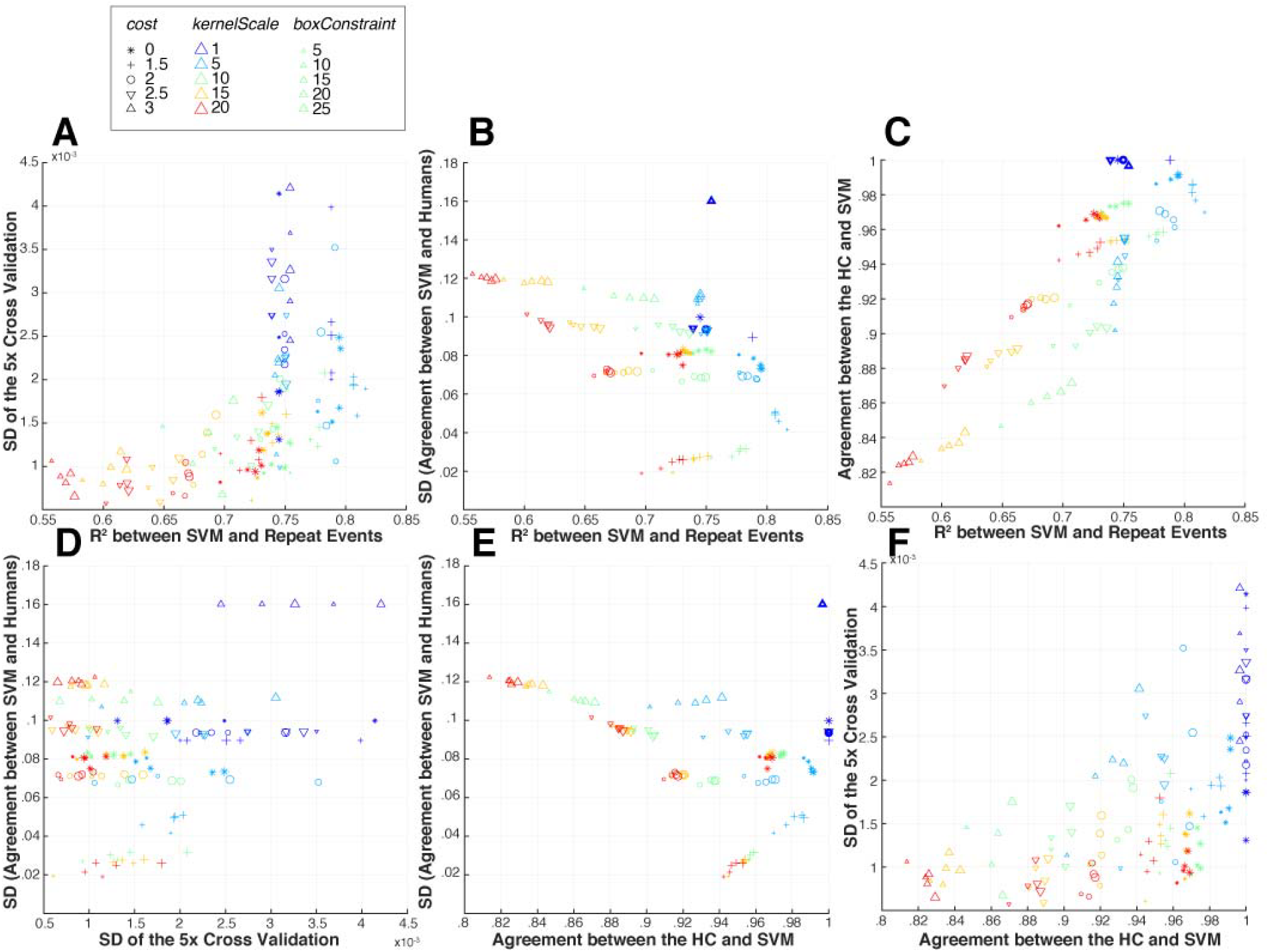
Parameter optimization for the SVM. We used a grid search method to vary box constraint, kernel scale, and the cost function parameters of the SVM. **(A-F)** Unique pair-wise combinations of our output parameters: 1) the standard deviation of the 5-fold cross fold validation, 2), the agreement between the humans and the SVM 3) the standard deviation of the agreement between the humans and the SVM, and 4) the R^2^ between the SVM classification score and the repeated event data set.

## References

1. McCormick DA, Contreras D. On the cellular and network bases of epileptic seizures. Annu Rev Physiol. 2001;63:815–46.

2. Huguenard JR, McCormick DA. Thalamic synchrony and dynamic regulation of global forebrain oscillations. Trends Neurosci. 2007;30:350–6.

3. Pinault D, Vergnes M, Marescaux C. Medium-voltage 5-9-Hz oscillations give rise to spike-and-wave discharges in a genetic model of absence epilepsy: In vivo dual extracellular recording of thalamic relay and reticular neurons. Neuroscience. 2001;105:181–201.

4. Sitnikova E, van Luijtelaar G. Electroencephalographic characterization of spike-wave discharges in cortex and thalamus in WAG/Rij rats. Epilepsia. 2007;48:2296–311.

5. Tan HO, Reid CA, Single FN, et al. Reduced cortical inhibition in a mouse model of familial childhood absence epilepsy. Proc Natl Acad Sci USA. 2007;104:17536–41.

6. Lowenstein DH. Seizures and Epilepsy. In Kasper DL (Ed) Harrison’s Principles of Internal Medicine. 19th Ed. New York, N.Y.: McGraw-Hill Education; 2015.

7. Boly M, Maganti R. Monitoring epilepsy in the intensive care unit: Current state of facts and potential interest of high density EEG. Brain Injury. 2014;28:1151–5.

8. Tzallas AT, Tsipouras MG, Tsalikakis DG, et al. Automated epileptic seizure detection methods: a review study. In Epilepsy-histological, electroencephalographic and phychological aspects 2012. InTech.

9. Webber WR, Litt B, Lesser RP, et al. Automatic EEG spike detection: what should the computer imitate? Electroencephalogr Clin Neurophysiol. 1993;87:364–73.

10. Fisher RS, Scharfman HE, deCurtis M. How can we identify ictal and interictal abnormal activity? In Issues in Clinical Epilpetology: A View from the Bench 2014 (pp. 3–23). Springer, Dordrecht.

11. Koutroumanidis M, Arzimanoglou A, Caraballo R, et al. The role of EEG in the diagnosis and classification of the epilepsy syndromes: a tool for clinical practice by the ILAE Neurophysiology Task Force (Part 1). Epileptic Disord. 2017;19:233–98.

12. Koutroumanidis M, Arzimanoglou A, Caraballo R, et al. The role of EEG in the diagnosis and classification of the epilepsy syndromes: a tool for clinical practice by the ILAE Neurophysiology Task Force (Part 2). Epileptic Disord. 2017;19:385–437.

13. Webber WRS, Lesser RP. Automated spike detection in EEG. Clin Neurophysiol. 2017;128:241–2.

14. Lüttjohann A. Disclosing hidden information in the electroencephalogram using advanced signal analytical techniques. J Physiol (Lond). 2017;595:7021–2.

15. Kornmeier J, Bach M. The Necker cube - an ambiguous figure disambiguated in early visual processing. Vision Res. 2005;45:955–60.

16. Richard CD, Tanenbaum A, Audit B, et al. SWDreader: A wavelet-based algorithm using spectral phase to characterize spike-wave morphological variation in genetic models of absence epilepsy. J Neurosci Meth. 2015;242:127–40.

17. Ovchinnikov A, Lüttjohann A, Hramov A, et al. An algorithm for real-time detection of spike-wave discharges in rodents. J Neurosci Meth. 2010;194:172–8.

18. Xanthopoulos P, Rebennack S, Liu C-C, et al. A Novel Wavelet Based Algorithm for Spike and Wave Detection in Absence Epilepsy. IEEE; 2010. pp. 14–9.

19. Bauquier SH, Lai A, Jiang JL, et al. Evaluation of an automated spike-and-wave complex detection algorithm in the EEG from a rat model of absence epilepsy. Neuroscience Bulletin. 2015;31:601–10.

20. Petrou S, Reid CA. The GABAAγ2(R43Q) mouse model of human genetic epilepsy. In Jespers Basic Mechanisms of the Epilepsies. 4^th^ ED. Bethesda (MD): National Center for Biotechnology Information (US); 2012.

21. National Research Council. Guide for the Care and Use of Laboratory Animals. National Academics Press. 2010.

22. Nelson AB, Faraguna U, Zoltan JT, et al. Sleep patterns and homeostatic mechanisms in adolescent mice. Brain Sci. Multidisciplinary Digital Publishing Institute; 2013;3:318–43.

23. Nelder JA, Mead R. A Simplex Method for Function Minimization. The Computer Journal. 1965;7:308–13.

24. Christianini N, Shawe-Taylor J. An Introduction to Support Vector Machines and Other Kernel-based Learning Methods. 2000.

25. Ben-Hur A, Weston J. A User’s Guide to Support Vector Machines. In: Data Mining Techniques for the Life Sciences. Totowa, NJ: Humana Press; 2009. pp. 223–39. (Methods in Molecular Biology; vol. 609).

26. Cawley GC, Talbot NLC. On Over-fitting in Model Selection and Subsequent Selection Bias in Performance Evaluation. Journal of Machine Learning Research. 2010;11:2079–107.

27. Muthukrishnan R, Rohini R. LASSO: A feature selection technique in predictive modeling for machine learning. IEEE; 2016. pp. 18–20.

28. Lloyd S. Least squares quantization in PCM. IEEE Transactions on Information Theory. 1982;28:129–37.

29. Maheshwari A, Akbar A, Wang M, et al. Persistent aberrant cortical phase-amplitude coupling following seizure treatment in absence epilepsy models. J Physiol (Lond). 2017;10:387–7260.

30. Halasz P, Terzano MG, Parrino L. Spike-wave discharge and the microstructure of sleep-wake continuum in idiopathic generalised epilepsy. Clin Neurophysiol. 2002;32:38–53.

31. Kelly KM. Spike–wave Discharges: Absence or Not, a Common Finding in Common Laboratory Rats. Epilepsy Currents. American Epilepsy Society; 2004;4:176–7.

32. Letts VA, Beyer BJ, Frankel WN. Hidden in plain sight: spike-wave discharges in mouse inbred strains. Genes, Brain and Behavior. Blackwell Publishing Ltd; 2014;13:519–26.

